# Membrane Topography-Driven Movement of Biomolecular Condensates

**DOI:** 10.1101/2025.03.24.644870

**Authors:** Matthias Pöllmann, Katja Zieske

## Abstract

Biomolecular condensates are assemblies of proteins or nucleic acids that exhibit liquid-like properties and organize intracellular biochemical reactions within many cells. Some condensates require membrane association, and we previously developed an assay to reconstitute biomolecular condensates in the presence of different membrane topographies. However, the effect of membrane topography on the displacement of biomolecular condensates remains incompletely understood. Here, we studied the movement of biomolecular condensates on lipid membrane-clad microstructures in a cell-free assay. We observed upward movements within microgrooves for untethered condensates. Increased membrane attachment reduced the number of upward movements. Further increasing the membrane attachment led to the formation of elongated condensates. We demonstrated a coordinated sideward movement of these elongated condensates. Finally, we found that molecular crowding with Ficoll70 decreased the frequency of upward movements by slowing condensate growth. Our results indicate that membrane topographies, in combination with membrane attachment patterns, regulate passive biomolecular condensate movement.

## INTRODUCTION

Biomolecular condensates are assemblies of proteins or nucleic acids within the intracellular space that regulate cellular function by compartmentalizing biochemical reactions.^1^ These condensates form through liquid-liquid phase separation, a process driven by intrinsically disordered domains or weak interactions of multivalent protein domains.^2,3^ Although biomolecular condensates are not enclosed by a lipid membrane and are thus also referred to as membrane-less organelles, various biomolecular condensates interact with cellular membranes.^4–6^ Upon signaling events such as phosphorylation, some condensates nucleate directly at membranes and remain membrane-associated .^2,7,8^ Others exhibit transient membrane binding.^9,10^ By interacting with lipid membranes, condensates regulate essential processes, including signal transduction and autophagy.^11–13^

The proper localization of biomolecular condensates is critical for cellular function^23^. One mechanism of condensate repositioning involves dissolution and re-condensation at specific cellular locations. This mechanism is important for P granules, which accumulate asymmetrically at the posterior of C. elegans germ cells to regulate asymmetric division.^10^ In contrast, RNA granules are actively transported over long distances along microtubules by hitchhiking on lysosomes, ensuring localized protein synthesis far from the nucleus.^9^ Lastly, Dishevelled proteins, after forming condensates in the cytosol, are recruited to the plasma membrane upon binding conformationally changed membrane proteins to initiate signaling cascades.^5,24^ While several mechanisms governing the movement of biomolecular condensates have been described, the full range of processes underlying the directional movement of biomolecular condensates remains to be elucidated.^25^

To investigate the physical principles governing condensate behavior, cell-free reconstitution experiments provide a powerful approach, offering precise control over condensate formation and composition. In addition, these assays provide opportunities to mimic condensate-membrane interactions through defined lipid compositions of model membranes^14–16^ and modulation of salinity or membrane charge^17^. Such variations have been used to study the wetting phenomena of biomolecular condensates. However, while salt concentrations affect not only the wetting of condensates, but simultaneously other characteristics,^18^ condensate-interacting lipids enable a more targeted approach for studying condensate wetting on lipid membranes.

Although the biochemical parameters governing the membrane recruitment of biomolecular condensates^2,17,19^ and the deformation of membranes by biomolecular condensates^16,20,21^ have been extensively studied, the influence of membrane topography itself is less well characterized. Most cell-free assays for studying the wetting of lipid membranes by biomolecular condensates use planar or spherical model membrane systems. However, cellular membranes feature protrusions and microdomains.^22^ Therefore, we previously developed an assay to reconstitute biomolecular condensates on topographically structured membranes and demonstrated that membrane topography affects the shape and localization of biomolecular condensates through capillary forces mediated by wetting and interfacial tension.^15^

Biomolecular condensates exhibit liquid-like properties, including fusion, fission, and molecular exchange with their surroundings and within the condensate.^10,26^ These properties suggest that principles from droplet physics may also apply to biomolecular condensates within cells. For example, the behavior of biomolecular condensates on structured lipid membranes may be similar to the behavior of water droplets on microstructured surfaces. In the presence of microstructured surfaces the motion of water droplets can be guided,^27^ with wetting properties influencing the droplet movement.^28^ The movement of water droplets, widely studied in material sciences and anti-fouling research, demonstrates how surface topography can influence liquid dynamics.^29,30^ We hypothesized that, similar to water droplets responding to surface topography, the topography of lipid membranes may direct the movement of biomolecular condensates.

Here, we use a cell-free assay to study the movement of biomolecular condensates on topographically structured lipid membranes. We show that condensates within membrane microgrooves move upwards, against gravity, once they have grown beyond a critical size. We demonstrate that this behavior primarily occurs in the absence of membrane interaction. Increasing the concentration of condensate-interacting lipids led to membrane wetting, causing condensates to remain preferentially within the microgrooves. Only larger condensates were able to perform upward movements within the microgrooves under these conditions. Even stronger membrane attachment resulted in the formation of elongated condensates, which exhibited a zipper-like sideward movement towards the grooves. This movement was induced by condensate fusion. Finally, we demonstrate that molecular crowding with Ficoll70 decreased the growth of protein condensates, thereby reducing the frequency of upward movements. Our observations indicate that lipid membrane topography affects the movement of biomolecular condensates. Wetting via protein-membrane interactions may thus act as a switch that directs biomolecular condensates into and or out of cellular membrane protrusions.

## RESULTS AND DISCUSSION

Membrane topography and mechanical confinement play essential roles in the coordination of cellular processes and the organization of biomolecular condensates. For instance, during cellular migration through small pores, diameter constrictions of a membrane-enclosed nuclear volume and chromatin deformations influences the shape and formation of biomolecular condensates.^31^ To study the effects of membrane topography on the dynamic localization of biomolecular condensates in a well-defined system and in the absence of additional cellular complexities, we employed a cell-free assay. Specifically, we reconstituted proteins that assemble into biomolecular condensates on microstructured PDMS surfaces coated with lipid membranes (Fig. 1A). Thereby, we selected an array of grooves as the topographical feature, because the bottom of these grooves represents an environment where protein condensates are confined by two walls and it covers a large proportion of the total area. Furthermore, condensates within the grooves can still grow through lateral fusion and enable facile analysis of individual biomolecular condensates. The PDMS microgrooves were fabricated using photolithography and soft molding. To visualize the surface topography and confirm the structural integrity of the microgrooves, we perfomed scanning electron microscopy (Fig. 1B). Lipid bilayers were generated via vesicle fusion, and laser scanning confocal imaging was usedto confirm the presence of continuous lipid membranes that adopted the topography of the PDMS microgroove topography (Fig. 1C). To induce the formation of biomolecular condensates, we added the two multivalent proteins PRM_4_ (40 µM) and SH3_4_-6xHis (40 µM). PRM_4_ consists of four identical proline-rich motifs (PRMs) of the protein ABL1. The protein SH3_4_-6xHis contains four identical SH3 domains derived from NCK1 (Fig. 1D)^2^ and an n-terminal histidine-tag. The histidine-tag facilitates membrane binding via interactions with the functional lipid DGS-NTA.

**Figure 1:**
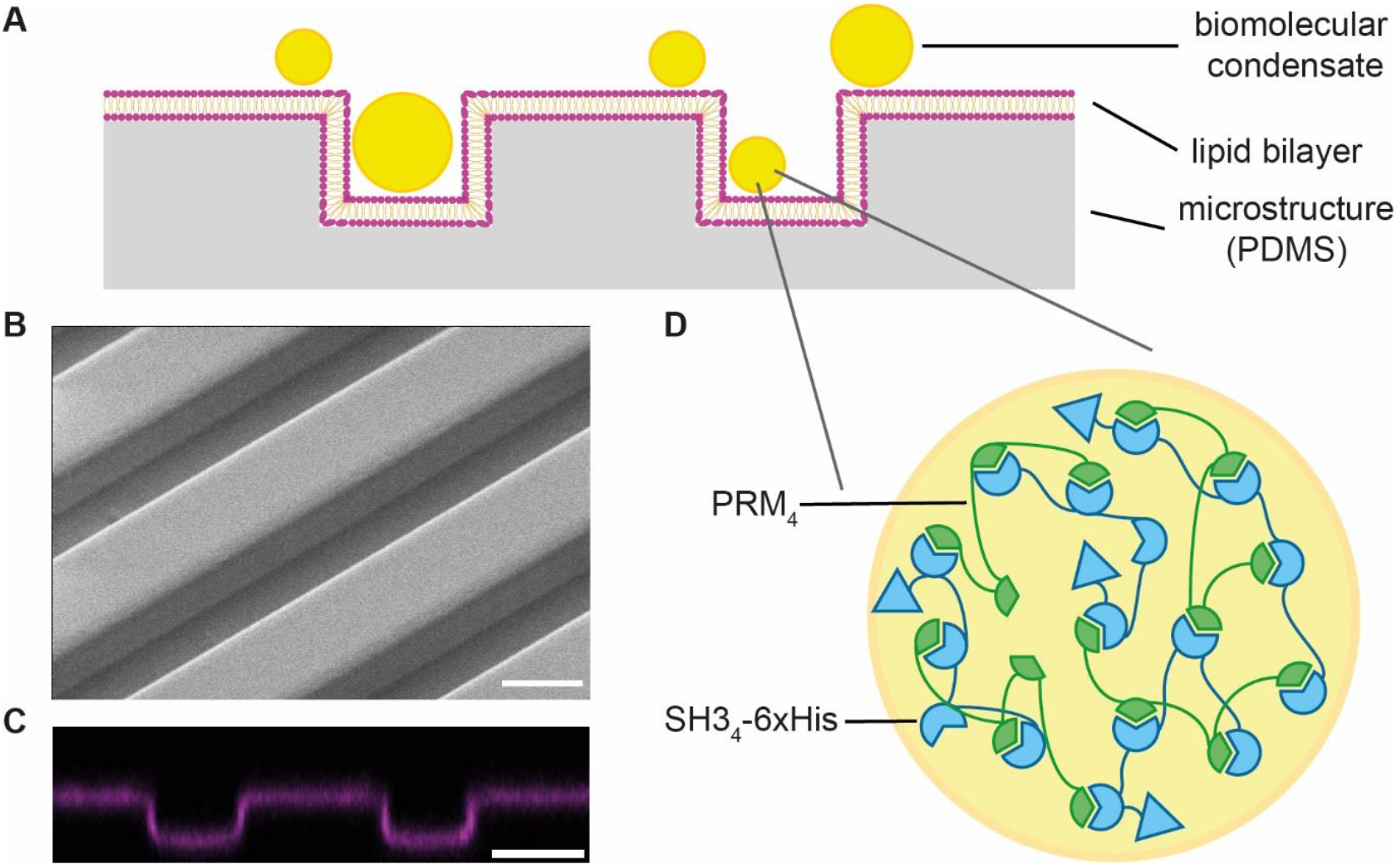
Cell-free assay for the characterization of biomolecular condensates on topographically structured membranes. (A) Schematic representation of the experimental system: Microstructured PDMS surfaces were clad with a lipid membrane. Liquid biomolecular condensates were assembled in the buffer solution above these lipid membranes. (B) Scanning electron microscopy image of microstructured PDMS surface. (C) Fluorescent laser scanning confocal microscopy image of a supported lipid bilayer labelled with 0.05 mol% Fast DiI. (D) Schematic illustration of biomolecular condensate composition. Biomolecular condensates were composed of the two proteins PRM_4_ and SH3_4_-6xHis. The SH3 domains interact weakly with PRM domains, and based on this interaction, the liquid biomolecular condensates were assembled. Scale bars: 10 µm.

To explore how confinement from two sides affects the dynamics of biomolecular condensates, we performed time-lapse confocal microscopy on biomolecular condensates within membrane-clad microgrooves. Thereby, we observed an interesting upward movement of large biomolecular condensates in the absence of membrane attachment (Fig. 2A, B). Initially, these condensates grew in the buffer solution through fusion and incorporation of protein monomers, and sedimented toward the microstructured bottom of the sample chamber, where the biomolecular condensates continued to grow. Condensates maintained a spherical morphology, and when their diameters exceeded the groove width, the condensates moved upwards and regained their spherical shape above the confining microgrooves.

**Figure 2:**
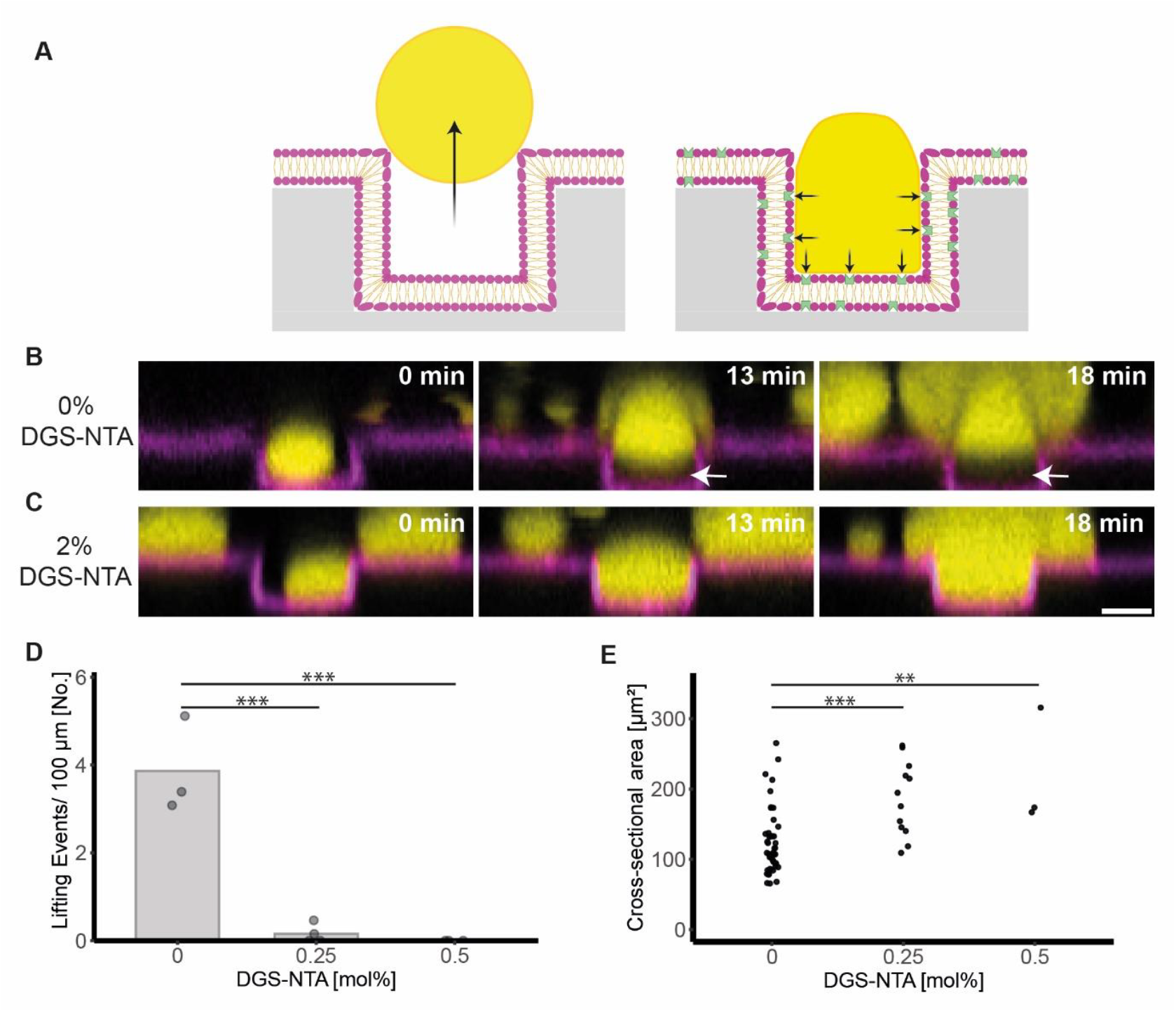
Upward movement of biomolecular condensates in microgrooves against gravity is determined by the absence of membrane attachment. (A) Schematic representation of biomolecular condensate dynamics within microstructured grooves in the absence (left: upward movement) and presence (right: remaining at the bottom of the microgrooves) of DGS-NTA – a membrane component that binds to the histidine tag of SH3_4_-6xHis. (B, C) Confocal time-lapse images of biomolecular condensates (yellow) within membrane grooves (magenta) in the absence (B) and presence (C) of 2 mol% DGS-NTA. The time-point of the first image of the depicted time-lapse sequences was defined as t=0. Independent samples for each condition: n=3. Scale bar: 5 µm. (D) The number of condensates displaying upward movements was quantified per 100 µm groove length and within a period of 35 min in the absence of DGS-NTA and in the presence of 0.25 mol% and 0.5 mol% DGS-NTA. Bar plot: Average number of biomolecular condensates displaying upward movements. Independent samples: n≥3. Two-tailed Student’s t-test: p=0.00026 (0% vs. 0.25% DGS-NTA) and p=0.00031 (0% vs. 0.5% DGS-NTA). (E) The cross-sectional areas of condensates after detachment from the groove bottom was determined. Cross-sectional area measurements were performed 3 µm above the top of the PDMS microstructures. Two-tailed student’s t-test: p = 0.0006 (0% vs. 0.25% DGS-NTA) and p=0.0051 (0% vs. 0.5% DGS-NTA). Number of measured condensates across three independent samples per condition: 47 (0% DGS-NTA), 12 (0.25% DGS-NTA), and 3 (0.5% DGS-NTA). Asterisks indicate statistical significance: p<0.05 (*), p<0.01 (**), and p<0.001 (***). Protein concentrations in all experiments: 40 µM PRM_4_, 40 µM SH3_4_-6xHis

This upward movement can be attributed to the interfacial tension of growing biomolecular condensates, which stabilizes their spherical shape. The system seeks to minimize the energetic cost of confinement by reducing contact with the groove walls, resulting in a displacement towards a less confined space. At the upper edges of the grooves, the absence of constriction reduces mechanical pressure at the droplet boundaries and this pressure gradient drives the protein condensates upwards. The observed upward movement of biomolecular condensates closely resembles the motion of water droplets within superhydrophobic grooves.^32^ Although such water droplets moved at air interfaces and involve larger assay, the similar behavior suggests that both systems may share the same underlying principles.

Next, we assessed how increased membrane wetting affects condensate dynamics. Our previous work demonstrated that DGS-NTA, which binds strongly to histidine-tagged condensate components, promotes membrane wetting by the condensates and reduces the membrane contact angle as compared to condensates on membranes without DGS-NTA.^15^ When we supplemented lipid membranes with 2 mol% DGS-NTA, upward movement of condensate was no longer observed (Fig. 2A, C). When we decreased the DGS-NTA concentration to 0.5 mol%, we still did not observe any upward movements within 35 minutes, indicating strong adhesion of the condensates to the membrane. When we decreased the DGS-NTA concentration further to 0.25 mol%, some condensates, which moved upward, were observed (Fig. 2D). However, without DGS-NTA the frequency at which upward moving condensates were observed, was significantly higher (Fig. 2D).

After longer incubation times of 45 min, larger condensates formed, and upward moving condensates were observed even in the presence of 0.5 mol% DGS-NTA. We measured the cross-sectional area of condensates after detachment from the bottom of the grooves (Fig. 2E), confirming that condensates moving upwards were on average smaller in the absence of DGS-NTA, as compared to upward moving condensates in the presence of DGS-NTA. Our data indicate that the size of condensates required for upward movements increases with increasing membrane attachment. This observation is in agreement with simulations of water droplets on microgrooves where wetting properties were modulated.^33^ Given that posttranslational modifications in cells can actively recruit proteins of biomolecular condensates to cellular membranes,^34^ cells may exploit wetting regulation as a switch to control condensate movement.

Next, we characterized how even stronger wetting influences the behavior of biomolecular condensates. For this, we supplemented the lipid membranes with 4 mol% DGS-NTA. Under these conditions, condensates in the grooves and those on the flat surfaces between them fused into bigger condensates, respectively. These fusions resulted in large, elongated condensates in the grooves and on the upper, flat surfaces between them (Fig. 3A, B). Initially, these condensates remained separate (Fig. 3B, left). However, when a condensate within a groove grew and reached the microstructure rim, it fused with adjacent condensates on the upper surface. In some instances, we observed fusion only on one side. Fusion was initiated at one contact point and then propagated along the length of the elongated biomolecular condensate (Fig. 3). The fusion process progressively pulled the corresponding condensates on the upper surface towards the condensate within groove, resembling a zipper-like motion along the microgrooves.

**Figure 3:**
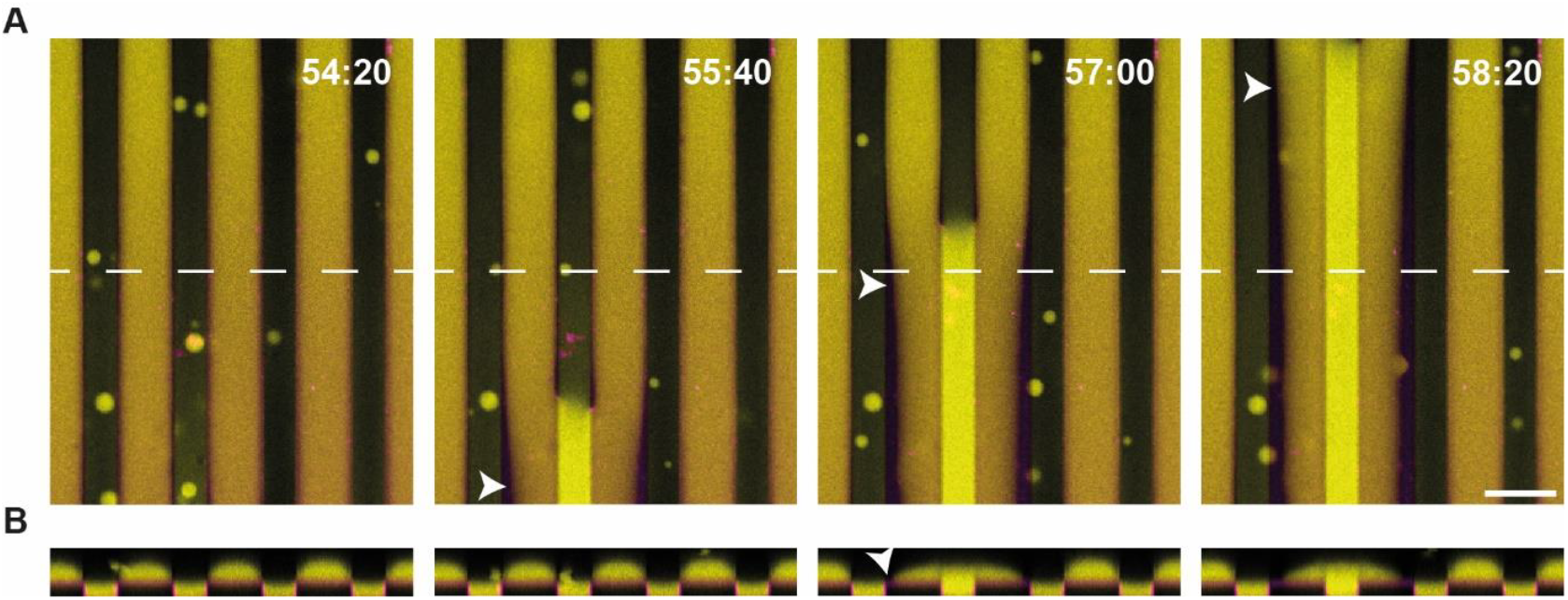
Lateral zipper-like movements of elongated biomolecular condensates on microgrooves are observed in the presence of 4% DGS-NTA. (A) Time-lapse confocal microscopy images of biomolecular condensates (yellow) in the presence of 4 mol% DGS-NTA. Elongated biomolecular condensates assembled within the valleys and at the upper level of membrane-clad PDMS surfaces with microgrooves (magenta). When condensates on the upper PDMS-level fused with condensates in the microgroove valleys, the condensates on the upper PDMS-level exhibited a lateral shift toward the microgroove. This resulted in a condensate-free area along the opposite edge of the upper PDMS level (white arrowheads). (B) Side view of biomolecular condensates on the same membrane surfaces along the white dotted line in (A). Protein concentrations: 40 µM PRM_4_ and 40 µM SH3_4_-6xHis. Independent samples: n=3. Scale bar: 20 µm.

The formation of elongated biomolecular condensates results from the interplay of microstructure, interfacial tension, and the strength of membrane attachment. We showed previously that other membrane-clad microstructures generate ordered patterns of biomolecular condensates.^15^ The present assay allowed us to generate elongated condensates and to study their planar zipper-like displacement. The observed coordinated movement may in the future provide insights into the mechanisms of physiologically relevant membrane-bound condensates, such as ZO-1 condensates in tight junctions.^35^

The intracellular environment is highly crowded, containing RNA, proteins, and other biomolecules. To determine how crowding affects the movement of biomolecular condensates, we mimicked molecular crowding using the crowding agent Ficoll 70. In the presence of 10% (w/v) Ficoll 70, we observed that proteins assembled into smaller condensates compared to non-crowded conditions (Fig. 4A, 4B). Our results show that molecular crowding with Ficoll 70 decreases the growth of PRM_4_/SH3_4_-6xHis condensates. We then examined how condensate size, modulated by crowding, affects upward movement. When we supplemented the buffer with Ficoll 70, fewer upward movements occurred within the microgrooves compared to non-crowded conditions, where condensates were larger (Fig. 4C). Previous studies on the related model proteins PRM_5_ and SH3_5_ demonstrated that Ficoll 70 acts as a volume-exclusion promoter, lowering the threshold concentration required for protein condensation.^36^ While this effect is well established, the reduced condensate growth observed in our setup may be caused by decreased diffusion and sedimentation in the crowded environment. Our results demonstrate that molecular crowding can be applied as a potent parameter in reconstitution approaches to generate and study smaller condensates. Since condensates assembled under crowding conditions were too small to exhibit upward movements within our microgrooves, our results further suggest that the relative size of condensates and membrane topography is important for driving upward movements of condensates.

**Figure 4:**
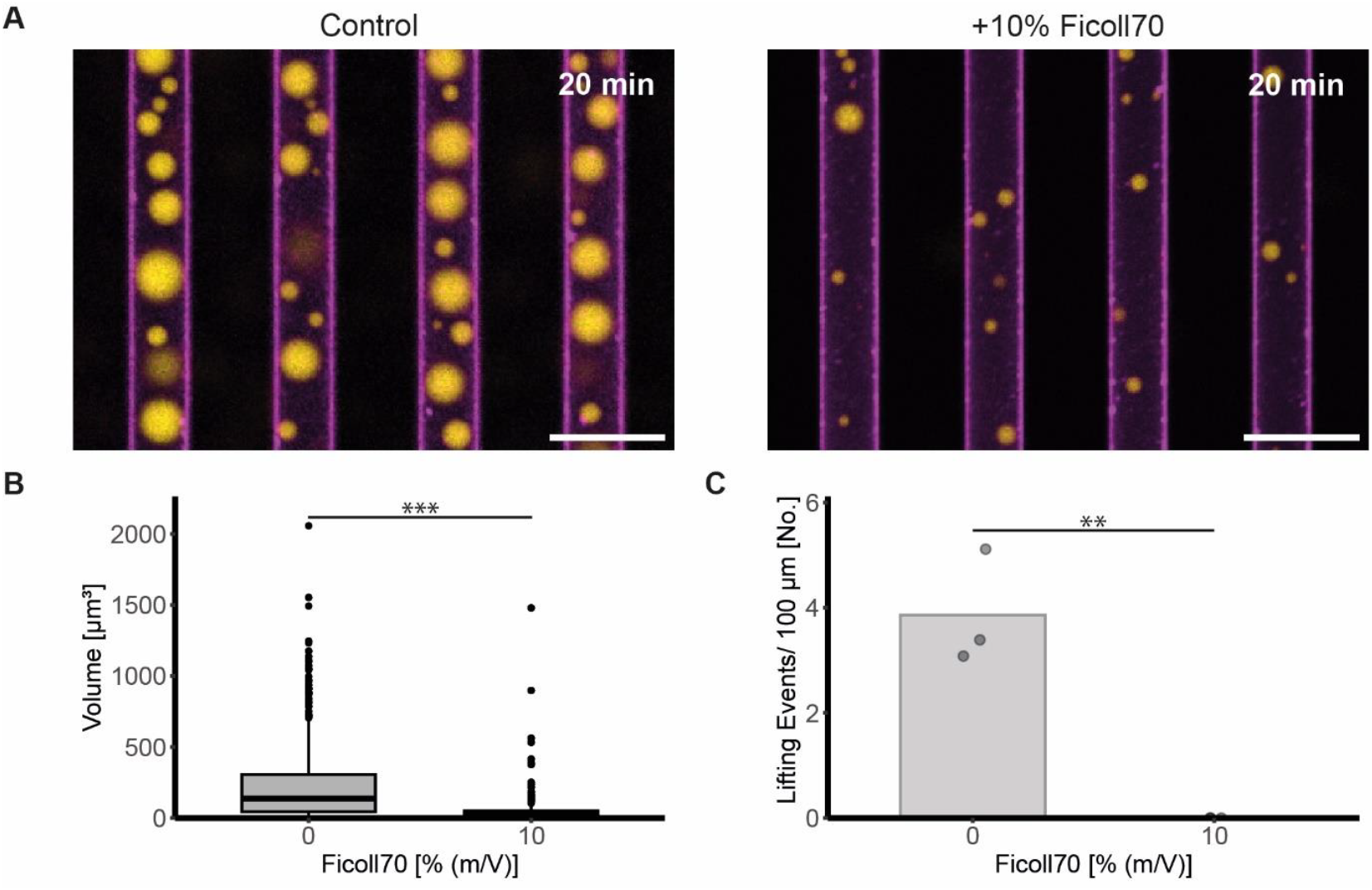
The number of condensates displaying upward movements within microgrooves is reduced, when condensates are small. (A) Confocal microscopy image of biomolecular condensates in microgrooves in the absence (left) and presence (right) of 10% (w/v) Ficoll 70. Protein concentrations: 40 µM SH3_4_-6xHis and 40 µM PRM_4_. Scale bar: 20 µm. Independent samples: n=3 (B) Volume measurement of biomolecular condensates after an assembly time of 20 min in the absence and presence of 10% (w/v) Ficoll 70. The volume of 441 (0%) and 256 (10%) biomolecular condensates larger than 0.5 µm^3^ was measured across three independent samples. Two-tailed Student’s t-test: p = <2e-16 (0% vs. 10% (w/v) Ficoll 70). (C) Number of biomolecular condensates displaying upward movements within 35 min and along a microgroove length of 100 µm in the absence and presence of 10% (w/v) Ficoll 70. Two-tailed Student’s t-test: p = 0.0036 (0% vs. 10% (w/v) Ficoll 70). Asterisks indicate statistical significance: p < 0.05 (*), p < 0.01 (**), and p < 0.001 (***).

In summary, we have shown that the motion of biomolecular condensates can be induced by interfacial tension and confinement. The behavior of biomolecular condensates in membrane-clad microgrooves thereby resembles the movement of water droplets on superhydrophobic microstructures and can be modulated by wetting of the condensates. Given the diverse topographical features of cellular membranes, these results may provide insights into the dynamic localization of biomolecular condensates within membrane-confined regions of the cell.

## MATERALS AND METHODS

### Cloning of plasmids

The plasmid encoding for SH3_4_-6xHis was generated by Gibson assembly from pGEX SH3(2)-4R (Addgene #112090).^2^ A high-fidelity PCR was performed using the following primers: 5’-tgtcatcatcatcatcatcactgaaggatccgcggccgcatcgt-3’ (forward) and 5’-cctcagtgatgatgatgatgatgacacaaaggactgtcaccttcttcagt-3’ (reverse). The amplified product was run on a 0.66% agarose gel. DNA was isolated from the band of the desired size. The linear DNA was annealed using the Gibson assembly cloning kit (New England Biolabs) and bacteria were transformed with this plasmid. Colony PCR and Sanger sequencing confirmed the correct sequence. The plasmid encoding for PRM_4_ (pMAL-Abl-PRM 4R, Addgene #112087) was previously mutated to introduce a cysteine tag for labeling.^15^

### Protein purification

Purification protocols were adapted from previously published protocols.^2^ Proteins were expressed in BL21(DE3) pLysS Competent Cells (Agilent Technologies, #200132). Transformed bacteria were grown at 37°C in Terrific Broth medium containing 0.1 mg/ml ampicillin. Protein expression was induced by adding 1 mM IPTG at an OD between 0.6 and 0.8. Proteins were expressed overnight at 18°C.

#### Purification of SH3_4_-6xHis

Bacteria were pelleted, resuspended in GST-Trap buffer A (pH=8.0, 50 mM Tris, 75 mM NaCl, 1 mM DTT) and lysed by sonication on ice. Lysates were cleared by centrifugation at 15,000g for 30 min at 4°C and incubated with Pierce™ Glutathion-Agarose beads (Thermo Scientific) at 4°C for at least 3 hours. Proteins were eluted using GST-Trap buffer B (pH=8.0, 50 mM Tris, 75 mM NaCl, 1 mM DTT, 25 mM reduced glutathione) and the GST-tag was cleaved with TEV protease at room temperature for 2-3 hours. The cleaved products were further purified using a HiResQ anion exchange column (Cytiva). The column was equilibrated, loaded with the protein solution, and washed with HiResQ buffer A (pH=8.0, 50 mM Tris, 1 mM DTT, 100 mM NaCl). Proteins were eluted by gradient elution over 30 column volumes with HiResQ buffer B (pH=8.0, 50 mM Tris, 1 mM DTT, 1 M NaCl).

#### Purification of PRM_4_

Bacteria were pelleted, resuspended in HiTrap buffer A (pH=7.5, 50 mM phosphate buffer, 100 mM NaCl, 20 mM imidazole) and lysed by sonication on ice. Lysates were cleared by centrifugation at 15,000g for 30 min at 4°C. Proteins were purified using a HiTrap Chelating HP column (Cytiva), and eluted in HiTrap B Buffer (pH=7.5, 50 mM phosphate buffer, 100 mM NaCl, 500 mM imidazole). Subsequently, the protein tags were cleaved by incubating the proteins with TEV protease overnight on ice. Cleaved products were separated on a HiResS cation exchange column (Cytiva). The column was washed with wash buffer (pH=7.5, 50 mM phosphate buffer, 1 mM DTT, 100mM NaCl) and gradient eluted over 10 column volumes using the elution buffer (pH=7.5, 50 mM phosphate buffer, 1 mM DTT, 1M NaCl). PRM_4_ with a cysteine tag was concentrated to 100 µM and labeled with Alexa Fluor™ C_5_. The unbound dye was removed using a PD-10 Desalting Column (Cytiva).

The buffer of the purified proteins was exchanged by dialysis in KMEI buffer (pH=7.5, 150 mM KCl, 1 mM MgCl_2_, 1 mM EGTA, 10 mM imidazole) at 4°C overnight. Protein solutions were concentrated to approximately 200 µM using a Vivaspin® (Sartorius). Aliquots were snap frozen in liquid nitrogen and thawed only once.

### Preparation of vesicles and supported lipid membranes

Supported lipid membranes were generated by vesicle fusion. The lipids DOPC and 18: 1 DGS-NTA (Ni) (dissolved in chloroform; Avanti Polar Lipids) and 0.05 mol% Fast DiI (dissolved in ethanol, ThermoFisher) were mixed in a glass vial. The lipid solution was dried under a nitrogen stream and then desiccated under vacuum to remove residual solvents. The lipid film was resuspended in membrane buffer (pH=7.5, 25 mM Tris, 150 mM KCl) to a concentration of 4 mg/ml, incubated at 37°C for 30 min and sonicated. The vesicle solution was aliquoted, frozen and thawed only once.

To generate supported lipid membranes, the vesicle solution was diluted to 0.5 mg/ml in membrane buffer, supplemented with 4 mM CaCl_2_ and added to a custom-made reaction chamber with a bottom consisting of a PDMS layer with microstructures. The lipid solution was then incubated at 37°C for 30 min and a lipid bilayer membrane assembled. The residual vesicles were removed by repeated dilution of the solution above the membrane. Finally, membrane buffer was replaced with KMEI buffer, or KMEI buffer supplemented with Ficoll 70.

### Preparation of PDMS structure

PDMS microstructures were prepared as described previously.^15^ The two-dimensional geometry of the microstructures was defined by patterns on a chrome mask. Approximately 7 µm high photoresist structures (mr-DWL 5, micro resist technology) were fabricated on silicon wafers by photolithography. The wafers were then treated with trimethylchlorosilane (Sigma-Aldrich). Microstructured PDMS surfaces were fabricated by soft molding. PDMS (Sylgard184, monomer to crosslinker ratio 10:1, Dow Corning) was degassed and poured on the silicon wafers. Glass coverslips were pressed through the liquid PDMS onto the microstructures. The PDMS was cured by incubation at 60°C for at least 3 hours. The glass coverslips with a layer of microstructured PDMS were carefully recovered. Surface structure was visualized using scanning electron microscopy (GeminiSEM 560, Zeiss). Before assembling lipid membranes, PDMS surfaces were treated with oxygen plasma at 0.3 mbar for 1 min to increase wettability and a plastic cylinder was glued on top of the PDMS surface.

### Microscopy

Microscopy images were acquired using a laser scanning fluorescence microscope (LSM980, Zeiss) with a 20x objective (Plan-Apochromat 20x/ 0.8, ∞/0.17, Zeiss). Microscopy experiments were performed at room temperature. Protein assays were performed in KMEI buffer. Protein condensation was induced by mixing 40 µM SH3_4_-6xHis and 40 µM PRM_4_. 5% of PRM_4_ was labelled with AF488. Time measurements started when both proteins had been added to the sample chamber, unless otherwise noted.

### Software

ImageJ/Fiji (v2.14.0/ 1.54f) was used for Image analysis.^37^ Volume measurements were conducted using 3D Objects Counter.^38^ The plug-in 3D Viewer was used to reconstruct 3D images displayed as volume from z-stack images.^39^

Subsequent data analysis and statistical tests were performed in R (v4.3.2) using RStudio (v2023.12.1). Adobe Illustrator (v28.7.1) and Blender (v4.0) were used to arrange figures and generate schematics.

## Acknowledgements

We thank all members of the Zieske research group for scientific discussions. The authors thank Eduard Butzen and the “Technology Development and Service Unit – Micro- and Nanostructuring” for the acquisition of scanning electron microscopy images. The plasmids pMAL-Abl-PRM 4R (Addgene plasmid #112087) and pGEX SH3(2)-4R (Addgene plasmid #112090) were gifts from Michael Rosen. Language and grammar of the manuscript was refined with assistance of Chat GPT (v3.5) and DeepL Write (March 2025).

This work was supported by a Max Planck Research Group grant from the Max Planck society awarded to Katja Zieske.

